# Closing the research-implementation gap using data science tools: a case study with pollinators of British Columbia

**DOI:** 10.1101/2020.10.30.362699

**Authors:** Laura Melissa Guzman, Tyler Kelly, Lora Morandin, Leithen M’Gonigle, Elizabeth Elle

## Abstract

A challenge in conservation is the gap between knowledge generated by researchers and the information being used to inform conservation practice. This gap, widely known as the research-implementation gap, can limit the effectiveness of conservation practice. One way to address this is to design conservation tools that are easy for practitioners to use. Here, we implement data science methods to develop a tool to aid in conservation of pollinators in British Columbia. Specifically, in collaboration with Pollinator Partnership Canada, we jointly develop an interactive web app, the goal of which is two-fold: (i) to allow end users to easily find and interact with the data collected by researchers on pollinators in British Columbia (prior to development of this app, data were buried in supplements from individual research publications) and (ii) employ up to date statistical tools in order to analyse phenological coverage of a set of plants. Previously, these tools required high programming competency in order to access. Our app provides an example of one way that we can make the products of academic research more accessible to conservation practitioners. We also provide the source code to allow other developers to develop similar apps suitable for their data.

## Introduction

The research-implementation gap is a well known phenomenon in conservation science, where over two-thirds of conservation assessments never lead to conservation action (Knight et al. 2008). This occurs when there is a mismatch between the information available for use by conservation practitioners and the information available in the scientific literature (Dubois et al. 2020). The responsibility of narrowing the research-implementation gap falls on both conservation practitioners and researchers and requires that they both improve communication in order to engage in effective conservation decision-making (Dubois et al. 2020).

Scientific knowledge is rarely shared and transferred effectively to the public and/or practitioners (Born et al. 2009). Scientists need to communicate with non-scientists in order to extend the findings of their research beyond peer-reviewed publications. One strategy is to actively distill and disseminate information — vs. passively disseminating information in peer reviewed journal articles. Actively disseminating information to a lay audience requires continued communication between researchers and conservation practitioners through activities such as meetings and working groups (Gibbons et al. 2008). Beyond active dissemination, researchers can engage stakeholders and policy makers in the process of developing conservation strategies and tools in order to meet the need of these stakeholders and policy makers — this is the goal of translational ecology (Wall et al. 2017).

Engaging stakeholders from the outset of a research project helps ensure that the resultant products meet their desired needs. For example, in a climate project in the Great Basin region of the US, resource managers collaborated with researchers to incorporate climate modelling into resource planning. One of the first hurdles they encountered was that staff did not have the expertise to apply technical research results (Wall et al. 2017). Because they had included these staff and their managers from an early stage in the project development, they were able to identify these problems and modify their research outputs to ensure they were usable by the staff.

Here, we design an interactive web app that builds the foundation for addressing this research-implementation gap for pollinator conservation in British Columbia in three ways: (i) actively disseminating information by collating data previously published in many manuscripts into a single database, (ii) lowering the technical barrier to entry by making the information easily accessible and manipulable, and (iii) deliberately engaging with conservation practitioners to produce an application of available data that would meet their needs. We partnered with Pollination Partnership Canada to ensure that the resultant web app would meet general and specific pollinator conservation needs.

### Conservation practitioner engagement

We worked with Dr. Lora Morandin, the Western Canada Program Director at Pollinator Partnership Canada — an NGO devoted to the conservation of pollinators— so that our tools could address their need for more information about pollinator distributions and plant associations, aligning with their goals to create and maintain habitat that supports native pollinator populations and pollinator diversity. One of their ongoing challenges is providing locally-relevant information to land managers on the potential impacts of conservation efforts (largely in the form of habitat restoration) for pollinators. While there is ample information on the effects of hedgerow restoration for pollinators in places like California (Kremen et al. 2004, 2019), past research conducted in British Columbia has been less well disseminated to practitioners. Our discussions identified the development of a web app providing information on plant-pollinator relationships would be an effective way to overcome this challenge. The app can also help conservation practitioners more generally by providing information about the plants most often visited by pollinators for pollen and nectar resources; such plants would be useful to include in habitat restorations focussed on supporting pollinator diversity.

### Data description

We collated data from 17 studies that documented some aspect of plant-pollinator communities across British Columbia. While the aim of each study was different, in all studies, pollinators were collected as they visited flowers. This type of data provides insight not only about the abundance and richness of pollinator communities, but also about the types of flowers that pollinators visit and how these visitation rates differ across regions of the province. This can be useful when designing planting schedules aimed at attracting pollinators to crops or for planning rural and urban flower strips for pollinator conservation. Due to the distribution across studies, these data are not currently available in a single, easily accessible format, despite being previously published across several separate manuscripts (Courcelles et al. 2013; Neame et al. 2013; Chamberlain et al. 2014a, 2014b; Wray et al. 2014; Gielens et al. 2014; Button & Elle 2014; Wray & Elle 2015; Elwell et al. 2016; Gibbs et al. 2016; Gillespie et al. 2017; Gillespie & Elle 2018; Toshack & Elle 2019; Kelly & Elle 2020a, 2020b). The collated dataset encompasses surveys of pollinators on plants from 2005 to 2017 across 196 individual sites (8 ecoregions from British Columbia). It contains over 30,000 records of pollinators and the plants they interact with. The data contains both native and non-native plants, and crops.

### Potential uses

#### Support of rare, at risk species or monitor introduced species

The western bumble bee (*Bombus occidentalis*) has declined substantially since 1998 (Graves et al. 2020). The main drivers of declines for the western bumble bee include pesticides and land-use change. Conservation of this endangered bumble bee requires not only addressing these main drivers of decline, but also creating or protecting suitable habitat Conservation practitioners can use our app to identify which plant species are known to be visited by *Bombus occidentalis* and, further, whether this pollinator changes flower resources in different regions. In addition, conservation practitioners can use our app to monitor introduced species such as *Bombus impatiens* by identifying the flower resources used by that species.

#### Planting option for non-crop uses

Cities are increasingly promoting and incorporating pollinator plantings into their operations and land management (Hall et al. 2017). Urban plantings designed for pollinators have been shown to increase pollinator abundance and diversity within city boundaries (Hofmann & Renner 2020). Urban planners and gardeners could use our app to identify plant mixes that can grow within a specific region and that also support a high abundance and diversity of pollinators.

#### Supporting crops

Flower strips and hedgerows have been shown to enhance pest control and increase crop pollination in adjacent fields. While the effect of the flower strip and hedgerows are dependent on the type of crop and the landscape context (Albrecht et al. 2020), our app could be used as a tool to identify plant mixes that share pollinators with the crop, following the method of M’Gonigle et al. (2017).

#### Education on plants and pollinators

Cities and urban centers have become important sites for conservation of pollinators (Hall et al. 2017). Pollinators in urban settings can provide inspiration for school programs that aim to increase scientific literacy (Saunders et al. 2018). In addition to introducing students to standard insect collection techniques, our app could be used to teach students about simple ecological concepts such as species interaction networks and plant-pollinator interactions.

### Application description

#### Main selections

The app allows users to filter the data by region. Here, regions are ecosections of British Columbia (Demarchi 1996). We used ecosections, as they represent small scale ecosystem differences in British Columbia. Currently, we include data from the following regions: Nanaimo Lowland, Leeward Island Mountains, Southern Gulf Islands, Southern Okanagan Basin, Fraser Lowland, Southern Pacific Ranges and the Okanagan Range. We also have data from some sites in the Pacific Northwest of the United States. The application provides an interactive map showing where these regions are located (Figure 1A). The default option includes all of the regions.

**Figure 1:**
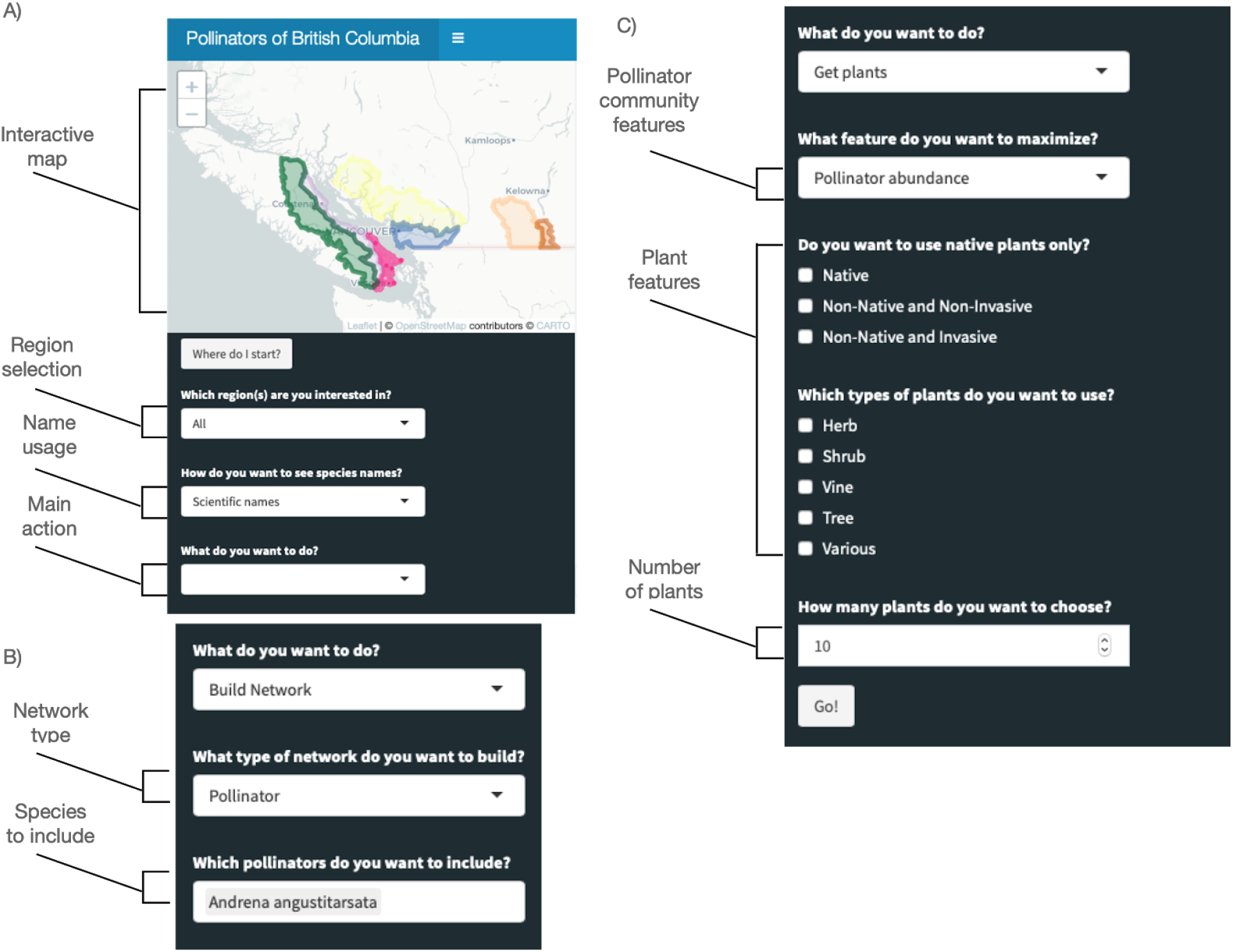
Layout and options in the application. A) Intro pane of application that allows the users to select the region, the names to be used and the main action. B) Options for building a network. C) Options for selecting groups of plants that support a crop or maximize pollinator abundance or richness.

Users can access the database using either scientific names (most of the specimens in the database are identified to species level) or common names, which typically correspond to genus level identification (Figure 1A). The default option is to use scientific names.

The app has three primary features: “Build network”, “Get Plants” and “Support Crop”. We will describe each of these features in detail below. These features allow users to visualize the data in three different ways and, in addition, help users select different plant mixes.

#### Build network

The “build network” feature allows the user to see all of the pollinators that visit a given plant or all of the plants that a pollinator visits. Therefore, this feature has two modalities, the “pollinator network” and the “plant network” (Figure 1B). The pollinator network allows the user to select pollinators and the resulting network plot shows the chosen pollinator along with the plants that the chosen pollinator visits. The plant network does the same as the pollinator network, but it allows the user to select plants. If a region is chosen, then the app will also filter the database to only show the plants or pollinators available for that region. The app will then produce a ‘network plot’, which shows the chosen plants/pollinators central to all of the species that they interact with (Figure 2). The thickness of the lines between plants and pollinators indicates the number of observations of that interaction. Finally a user can click on any of the species in the network and they will be directed to the Wikipedia page for that species.

**Figure 2:**
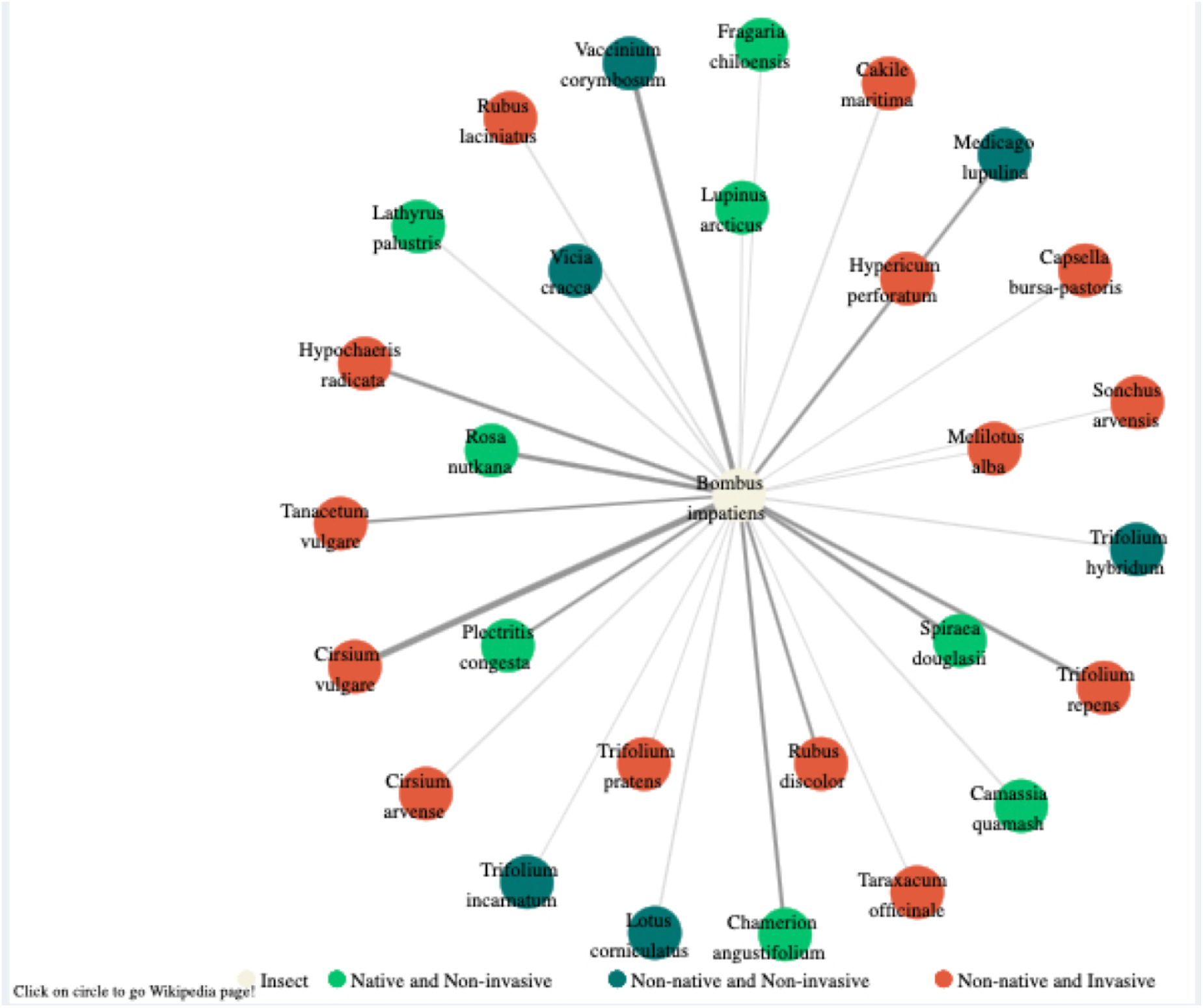
Example plots for the ‘Build Network’, using all the regions and *Bombus Impatiens* as the chosen pollinator.

#### Get plants

The “get plant” main option allows the user to select a specified number of plants that will be chosen to maximize either pollinator abundance, pollinator richness, or pollinator phenological coverage (Figure 1C). We refer to phenological cover as the weeks where a set of plants is flowering. In order to maximize pollinator abundance or richness, we filter the database based on the region and the plant attributes (such as whether the plants are native or non-invasive) and then sum the number of pollinator specimens or species, respectively, found on a given plant species (Figure 3).

**Figure 3:**
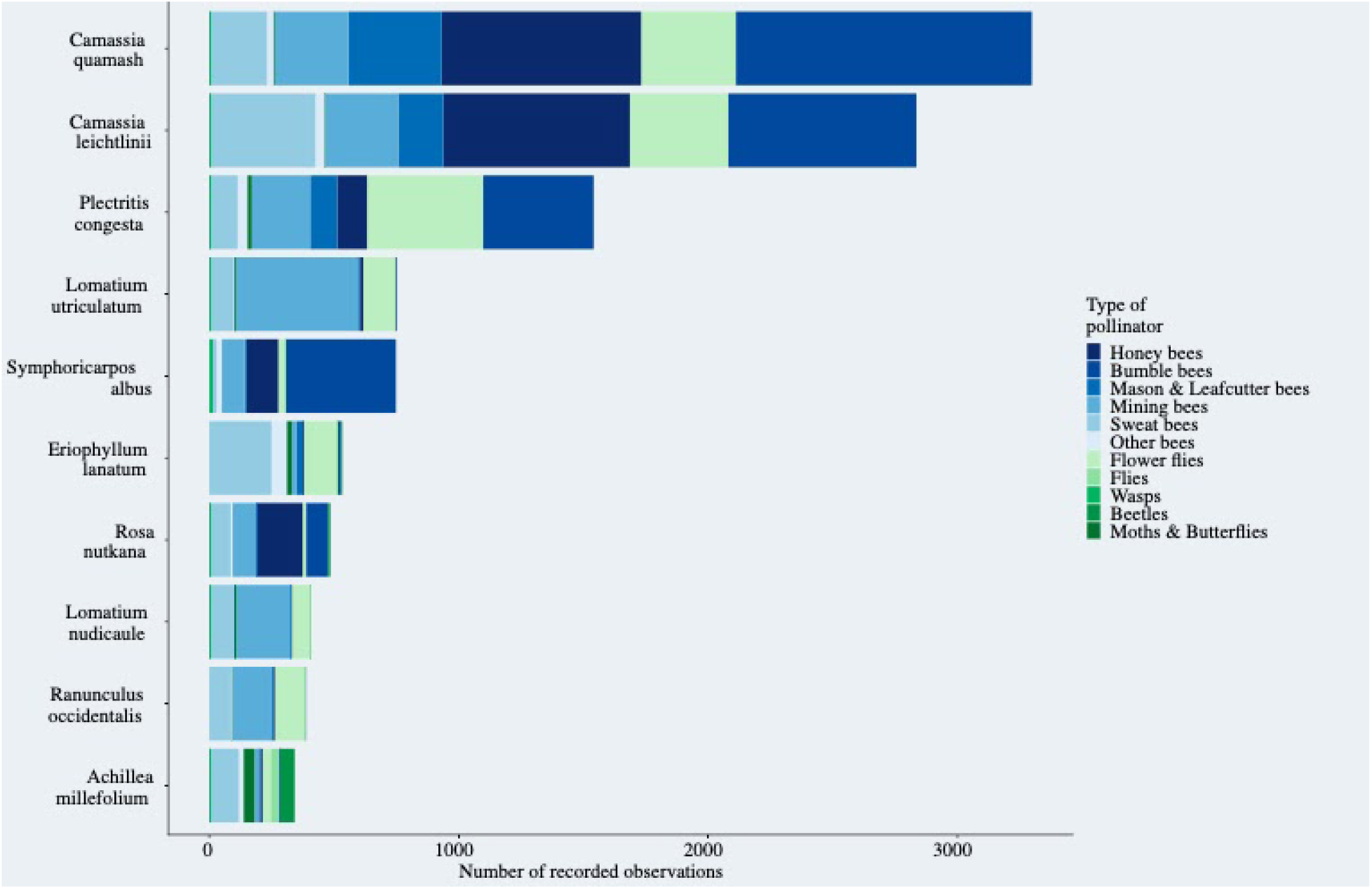
Example plot for the ‘Get plants’ feature. Here we selected to maximize pollinator abundance is all regions.

In order to maximize plant phenological coverage we use a genetic algorithm as developed in (M’Gonigle et al. 2017). Here phenological coverage is the number of weeks that a mix of plants is blooming. The genetic algorithm then finds the set of plants that minimize gaps in bloom periods. To run the algorithm we need the blooming period for each plant in the database. In order to obtain bloom periods for plants, we identify the weeks that each plant was observed to be ‘interacting’ with pollinators. While this is not an independent measure of blooming times for each plant, it is consistent across all of the different surveys done. Given that some of the observations may be outliers, and we want to focus mainly on ‘peak flowering times’, we removed the extreme 10th percent dates. Before running the algorithm we filter the database to the specified region and plant attributes. We also included an option for the user to consider only plants that bloom within a specified range of dates.

#### Support crop

Supporting a crop essentially uses the same algorithms and filters, but aims to maximize phenological coverage using the ‘Get plants’ option under the constraint that only pollinators that are known to visit the specified crop(s) are considered (Figure 4).

**Figure 4:**
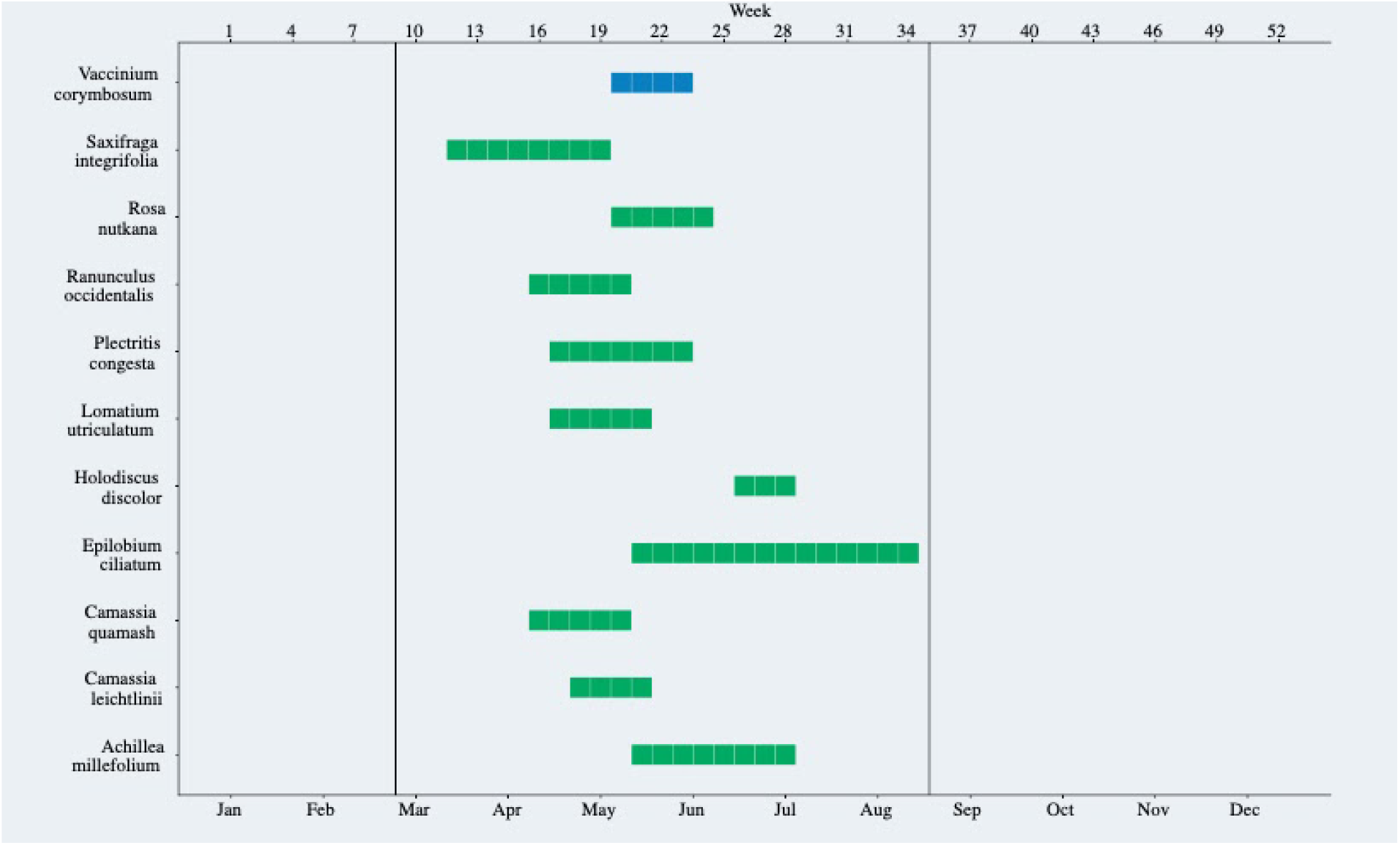
Example of the “Support crop” feature. In blue we show the crop (Blueberry) and in green the plant mix that would produce optimal phenological coverage for pollinators that are present during blueberry bloom.

One additional option that must be specified when selecting plants to support a crop is whether or not the selected plants can bloom at the same time as the crop. We included this option because allowing the support plants to bloom at the same time as the crop might ‘pull’ the pollinators away from the crop, but on the other hand it can also provide a more diverse diet to crop pollinators, and floral diversity inadjacent fields are known to improve crop pollination (Long et al. 1998; Garibaldi et al. 2013). Including this option allows users to tailor the plant mix to their needs. If the option selected is ‘Yes’, then the algorithm is the same as in ‘Get plants’. If the option selected is ‘No’, then we force the algorithm to contain the crop in every step so that the optimization of the flowering times is around the crop. While the algorithm can still select plants that overlap, the plants will be less likely to overlap with the crop.

The application was developed using the shiny package (Chang et al. 2017) in the R programming language (R core team 2019). All of the code to develop the application can be found in https://github.com/lmguzman/bcbees. The application is hosted at https://shiny.rcg.sfu.ca/bc-bees/.

## Caveats of the application

We worked closely with Pollination Partnership to ensure that the main options for the application would be useful for conservation practitioners wanting to support pollinator populations through habitat creation and maintenance. A major goal in design was to ensure that the application would be accessible to users who do not want or are not comfortable working with the raw data. This app is intended to provide only one tool when choosing plant species. The app does not include information about the commercial availability of plant species, or growing factors such as soil, moisture, and sun requirements. However, the plant growth requirements and preferences can be found online in numerous places. Assessing and incorporating plant availability is difficult to include, as information is limited, regionally variable, and not static. Users can generate plant lists that include more plants than they would use, ensuring that they will be able to obtain some of the species of interest. Finally, our app is limited by availability of data. While the datasets we have incorporated so far are extensive, additional data, particularly from natural habitats, would be beneficial. Currently, the available data was collected with an emphasis mostly on pollination in or around a single crop, Blueberry. The incorporation of more data is a major future goal.

## Future developments

This application is an evolving project and will continue to change as we get feedback from users. A limitation of the application is the quantity and quality of data; however, this is a living application and we intend to incorporate more data, first by expanding the geographic range and second by incorporating other sources of data such as citizen science data (Kremen et al. 2011). Finally, since this application was designed to suit the needs of end users, we will heavily rely on feedback from users for the next development stages.

## Conclusion

We present an interactive application and all of the source code used to develop it, with the goal of reducing the research-implementation gap in pollination conservation using data from British Columbia as a test of concept and first publicly-available tool. One of the main components of our approach was the co-development of the application with stakeholders who will use the application. This application can be used as a template to actively disseminate and make ecological research useful to different stakeholders.

## Acknowledgements

We would like to thank Melissa Platsko for compiling species links, Sarah Johnson, Elijah Reyes, Claire Kremen, Carly McGregor and the Native bee society of British Columbia for in-depth feedback for the shiny app. We acknowledge funding from Simon Fraser University (to LKM and LMG) and the Natural Sciences and Engineering Research Council of Canada (NSERC) (Discovery Grants to EE and LKM). The shiny app is hosted in the Simon Fraser University server.

